# Penalized Decomposition Using Residuals (PeDecURe) for Nuisance Variable Adjustment in Multivariate Pattern Analysis

**DOI:** 10.1101/2022.01.27.477859

**Authors:** Sarah M. Weinstein, Christos Davatzikos, Jimit Doshi, Kristin A. Linn, Russell T. Shinohara, Alzheimer’s Disease Neuroimaging Initiative

## Abstract

In neuroimaging studies, multivariate methods provide a framework for studying associations between complex patterns distributed throughout the brain and neurological, psychiatric, and behavioral phenotypes. However, mitigating the influence of nuisance variables, such as confounders, remains a critical challenge in multivariate pattern analysis (MVPA). In studies of Alzheimer’s Disease, for example, imbalance in disease rates across age and sex may make it difficult to distinguish between structural patterns in the brain (as measured by neuroimaging scans) attributable to disease progression and those characteristic of typical human aging or sex differences. Concerningly, when not properly adjusted for, nuisance variables can obscure interpretations and preclude the generalizability of findings from neuroimaging studies. Motivated by this critical issue, in this work we examine the impact of nuisance variables on features extracted from image decomposition methods and propose Penalized Decomposition Using Residuals (PeDecURe), a new MVPA method for obtaining nuisance variable-adjusted features. PeDecURe estimates primary directions of variation which maximize covariance between residualized imaging features and a variable of interest (e.g., Alzheimer’s diagnosis) while simultaneously mitigating the influence of nuisance variation through a penalty on the covariance between residualized imaging features and those variables. Using features estimated using PeDecURe’s first direction of variation, we train an accurate and generalizable predictive model, as evidenced by its robustness in testing samples with different underlying nuisance variable distributions. We compare PeDecURe to commonly used decomposition methods (principal component analysis (PCA) and partial least squares) as well as a confounder-adjusted variation of PCA. We find that features derived from PeDecURe offer greater accuracy and generalizability and lower partial correlations with nuisance variables compared with the other methods. While PeDecURe is primarily motivated by MVPA in the context of neuroimaging, it is broadly applicable to datasets where the dimensionality or complexity of the covariance structure calls for novel methods to handle sources of nuisance variation.

## 1 Introduction

The incorporation of neuroimaging data in research and clinical practice has brought new insights into how structural and functional patterns in the brain are associated with behavioral, psychiatric, and neurological phenotypes. For instance, structural magnetic resonance imaging (MRI) has given us a more nuanced understanding of changes in the brain associated with pre-symptomatic markers and longitudinal progression of Alzheimer’s Disease (AD) (Frisoni et al. 2010; Ferreira and Busatto 2011). However, since neuroimaging studies of AD and other disorders are often observational by design, controlling the impact of confounders and other nuisance variables is necessary for generalizability and interpretablity (Linn et al. 2016; Scheinost et al. 2019; More et al. 2020).

A standard approach to analyzing neuroimaging data is to use “mass-univariate” methods by modeling measurements independently at each location in an image and then synthesizing results across locations for interpretation. A mass-univariate approach was involved in some of the most foundational methods in neuroimaging research, including voxel-based morphometry (Ashburner and Friston 2000; Davatzikos et al. 2001) and statistical parametric mapping (Friston et al. 1991; Friston et al. 1994). Nuisance variable adjustment in the mass-univariate framework is also applied at the level of each imaging unit. One approach is to “regress out” nuisance variables like age and sex by fitting a separate regression model at each location in the image across subjects and subtracting out the effects of confounding variables at each location. Then subsequent analyses are conducted by combining these residualized features and estimating associations between those adjusted features and an outcome of interest (e.g., AD diagnosis). Another approach is to simply adjust for nuisance variables in location-specific regression models—for example, fitting a logistic regression model with AD diagnosis as an outcome and features of interest (e.g., regional brain volumes) and nuisance variables (e.g., age and sex) as covariates. In the mass-univariate framework, we would then visualize a brain map of test statistics derived from these location-specific models and interpret regions (or clusters of regions) whose estimated statistics exceed some threshold to be important in driving associations between brain structure or function and disease.

While mass-univariate methods continue to be widely used in neuroimaging research, is also understood that these methods fail to account for the complex spatial structure of the brain. Methods that do take this structure into account — often referred to as multivariate pattern analysis (MVPA) — are therefore increasingly preferred (Habeck et al. 2008; Habeck and Stern 2010; Westman et al. 2011). In this paper, we advocate for a multivariate approach to confounding adjustment in neuroimaging studies. To our knowledge, limited prior work has been done to adjust for confounders and other nuisance variables in the context of MVPA. One example is work by Linn et al. (2016), who weight the slack variables in the support vector machine objective function using inverse probability weights for a binary variable (e.g., disease status), more commonly used in confounding adjustment for treatment effects in epidemiological studies (Hernán and Robins 2006; Cole and Hernán 2008). Another example is Rao et al. (2017)’s instance weighting approach, which allows for weighting involving continuous outcome variables (e.g., clinical risk scores).

There is also a growing body of relevant work in nuisance variable adjustment of moderate-to highdimensional data outside the neuroimaging literature, including work in feature extraction and dimension reduction. Lin et al. (2016) propose adding a penalty term to the standard PCA objective function to achieve simultaneous batch effect correction across donors in a human brain exon array dataset. Aliverti et al. (2021) propose a similar penalized decomposition approach as Lin et al. (2016), aiming to remove undesirable correlations in predictive models, which may introduce unfair biases into predictive models. Tabak et al. (2020) use a domain adaptation framework for nuisance variable adjustment in cellular imaging. There is also some work using adversarial learning for the removal of confounding variables (Adeli et al. 2019; Zhao et al. 2020), highlighting the importance of confounding adjustment to achieve generalizability. While these methods vary in their applications, they share a similar goal, which is to correct for nuisance variables by mapping data to a space that is free from confounders, while also taking into account the inherently complex and correlated structure of the data. Subsequent analyses or predictive modeling may be conducted using this adjusted set of features.

In the present work, we examine the challenge of feature decomposition when goal is to both adjust for nuisance variables in complex data (e.g., brain images) and also preserve sources of variation in the data that are related to patterns of interest (e.g., those related to disease). To address these goals, we propose a new method: Penalized Decomposition Using Residuals (PeDecURe). While PeDecURe is primarily motivated by methodological issues in neuroimaging research, these issues are not unique to neuroimaging and the method can be easily applied to data across a variety of areas.

## 2 Methods

### 2.1 Notation and Previous Methods

Let *X* be an *n* × *p* matrix of *p* image-derived features, such as volumes of different regions of interest (ROIs) in the brain, for *n* study subjects. Without loss of generality, we assume each column of *X* has been centered to have mean zero. Let *A* be an *n* × *q* matrix of nuisance variables, which (on the basis of subject area knowledge) are considered important to account for before we can study the direct relationship between *X* and *Y*, an outcome of interest (say, AD diagnosis). *X*, *A*, and *Y* are assumed to have some degree of interdependence, which we illustrate in Figure 1(a). While *A* may include confounders as defined in a traditional causal inference framework (Greenland et al. 1999), in this work we use the term “nuisance variables” to refer more broadly to sources of variation that are not of direct interest, but which have some interdependence with variables that are of interest.

**Figure 1:**
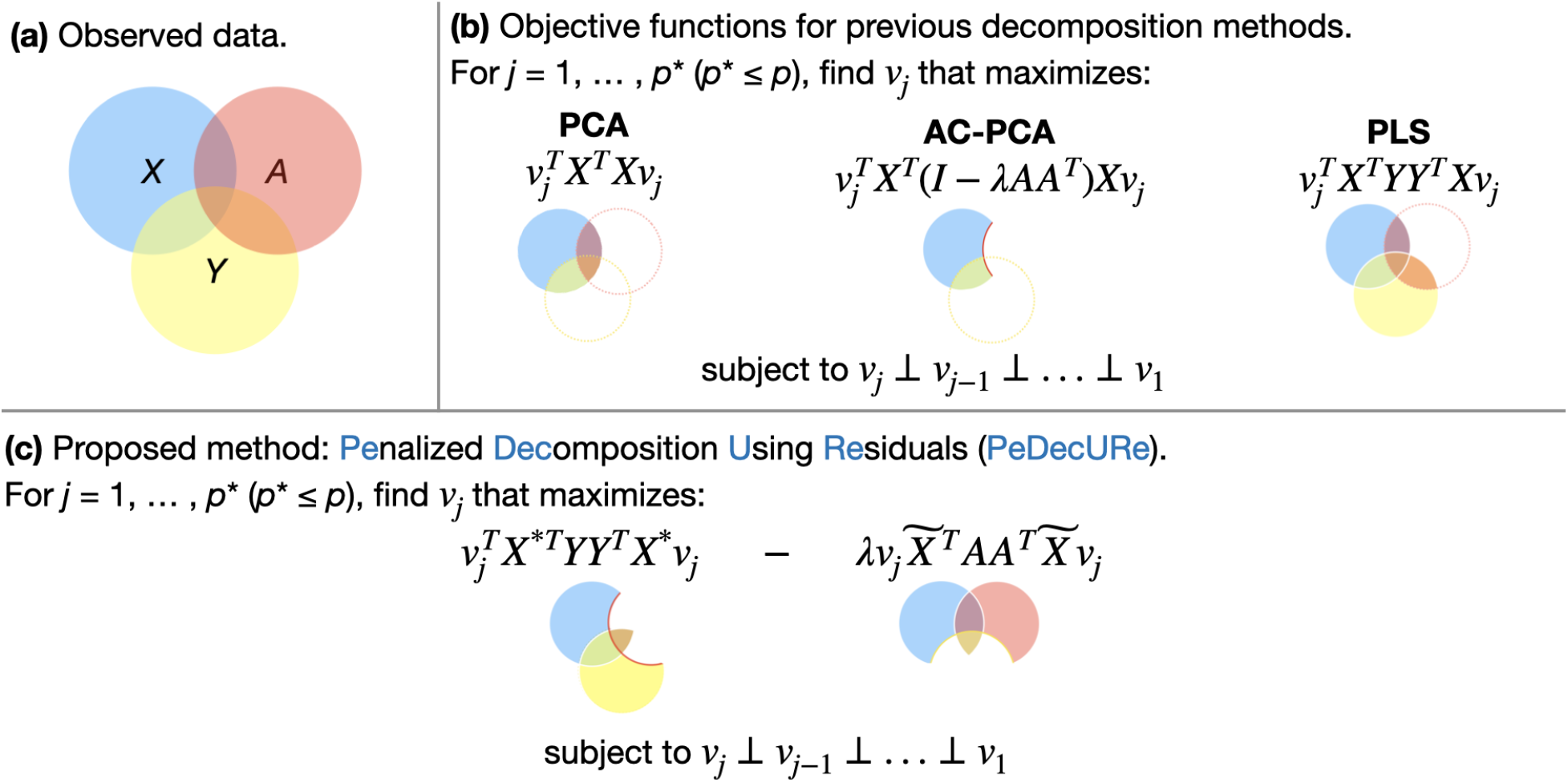
Visualization of observed data structure and objective functions for decomposition methods. In (b), we present objective functions for three existing methods and visualize the portion of the data whose covariance will be explained by *v_j_*. PCA solves *v_j_* that maximizes covariance of *X*, AC-PCA solves a similar maximization, but in a subspace that is orthogonal to (or approaches orthogonality to) *A* (as controlled by λ). PLS maximizes the covariance of *X^T^Y*, prioritizing shared variation between *X* and *Y*. In (c), we separately illustrate the portions of the data whose covariance we want to simultaneously maximize and penalize. We want *v_j_* that simultaneously maximizes associations between *X* and *Y*—but not the portions of *X* that are also associated with *A* (hence using the residual *X*\* instead of the original *X*))—and we also want *v_j_* to minimally explain covariation between *X* and *A*—without penalizing portions of *X* that are also associated with *Y* (hence the residual 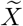).

Using this notation, the objective functions of PCA, partial least squares (PLS, in the case of a univariate outcome), and PCA with adjustment for confounders (AC-PCA, (Lin et al. 2016), although their method may also be more broadly applied to nuisance variables) are illustrated and provided in Figure 1(b). Briefly, each decomposition method estimates *v_j_* (*j* = 1, …, *p*\* for *p*\* ≤ *p*), which are directions of variation or primary components (PCs) that maximize the covariance of *X* (PCA) or covariance of *X^T^Y* (PLS), subject to orthogonality to previously estimated PCs (as well as other potential constraints).

We refer to the method that maximizes the covariance of *X^T^Y* as PLS, although the scenario involving a univariate outcome variable *Y* is only a special case of PLS. The eigendecomposition formulation of PLS differs from PCA in that it maximizes the covariance of *X^T^Y*, so the directions of variation that we obtain for *X* prioritize sources of variation that are shared between *X* and *Y*. In Figure 1(b), we illustrate the portion of the observed data whose covariance we aim to maximize in PCA (*X* (blue), including regions of overlap between *X* and other variables) and PLS (regions of overlap between blue (*X*) and yellow (*Y*)).

Recognizing that some part of the covariances of *X* and *X^T^Y* are shared with *A*, Lin et al. (2016)’s AC-PCA maximizes covariance of *X* in a restricted subspace with reduced confounding (in Figure 1, regions of *X* (blue) that have no overlap with *A* (red)), as determined by a tuning parameter, λ. Lin et al. (2016) accomplish this by appending a penalty term to the objective function of PCA, which penalizes covariance between *X* and *A*. However, as illustrated in Figure 1(b), the subspace of *X* that is maximized in AC-PCA may also remove certain information about *Y*, which is of interest.

### 2.2 Proposed Method: Penalized Decomposition Using Residuals (PeDecURe)

An important aspect of our proposed method for nuisance variable-adjusted feature decomposition is residualization of imaging features. We obtain these residuals through a standard mass-univariate approach. First, we fit the following linear model at each image location (although other more flexible or nonlinear models may also be considered at this stage):

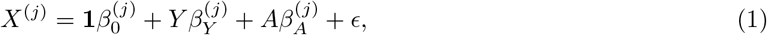

where where *X*^(*j*)^ is an *n*-dimensional vector of image measurements for each subject at locations *j* = 1, …, *p*, **1** is an *n*-dimensional vector of 1’s, 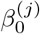 is the location-specific intercept parameter, *Y* is an *n*-dimensional vector of a variable of interest with one measurement per subject, 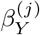 represents the effect of *Y* on the *j*th image location, *A* is the *n* × *q* matrix of nuisance variables, 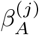 is a *q* × 1 vector of the effects of each column of *A* on the *j*th image location, and *ϵ* is an error term assumed to follow a standard normal distribution.

Parameter estimates from each location-specific model are used to compute the following residuals:

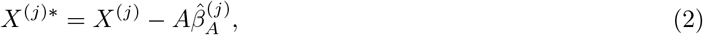

or the residuals that remais after regressing *A* out from the *j*th column of *X*, conditional on *Y*, and 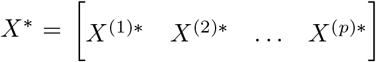 is the matrix of these residuals. We also define the residual that remains after regressing *Y* out from the *j*th column of *X*, conditional on *A*:

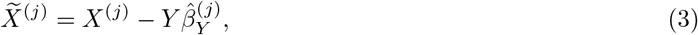

and 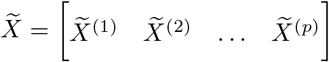 is the matrix of these residuals.

Each set of residuals plays a role in the formulation of PeDecURe’s objective function. Specifically, for *j* ≤ *p*, we solve:

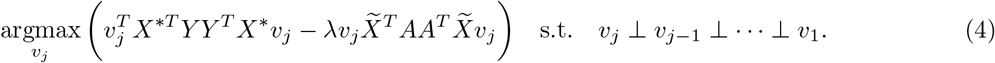

In words, PeDecURe identifies primary components (PCs) which maximize covariance between *X*\* and *Y*, while simultaneously penalizing associations between 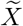 and *A* (also illustrated in Figure 1(c)). By incorporating different residuals in our maximization and penalty terms we reduce the potential for the PCs to be correlated with *A* or have attenuated correlation with *Y*.

Like in AC-PCA, PeDecURe’s penalization term is scaled by a tuning parameter, λ. In AC-PCA, λ controls the degree to which associations between features *X* and *A* are penalized and selected to maximize the ratio of covariance of *X* explained to the covariance between *X* and *A* explained (Lin et al. 2016). PeDecURe’s λ is rooted in a similar idea, but since we replace *X* with 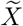 in the penalty term, we limit the scope of the penalization to the portion of shared variation between *X* and *A* that does not also overlap with *Y*. In each iteration of PeDecURe, we select a λ which, on average (across nuisance variables and PCs), maximizes the difference between the absolute value of estimated partial correlations between PC scores with *Y* (conditional on each *A*) and the absolute value of partial correlations between scores with each *A* (conditional on *Y*). Out of a set of user-pre-specified candidate, we use the following maximization to identify an optimal λ for a given dataset:

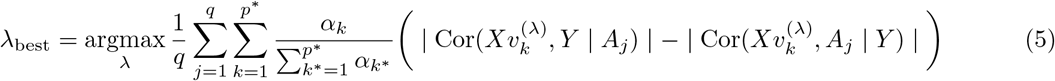

where *q* is the number of nuisance variables, *p*\* is the number of PCs being estimated, *α_k_* is the eigenvalue corresponding to the *k*th PC (so that 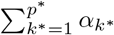 is the denominator for the portion of variation explained by *α_k_* when *p*\* PCs are being estimated), and 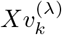 is the *k*th PC score from PeDecURe, estimated using tuning parameter value set to λ. We use the proportion of variation explained by each PC (ratio of each eigenvalue *α_k_, k* = 1, …, *p*\*, to the sum of eigenvalues through *p*\*) to up-weight earlier components in this maximization.

PC scores for PeDecURe and other methods considered in this paper (Figure 1) are computed in the same way: using the (column-centered) training and testing sample subsets of the feature matrix (denoted by *X_train_* and *X_test_*) we calculate the *j*th PC scores in the training and testing samples (using *v_j_* (*j* = 1, …, 3) estimated in each training sample) as PC_*j*(*train*)_ = *X_train_v_j_* and PC_*j*(*test*)_ = *X_test_v_j_*. Note that for all methods, we use the same *X_train_* and *X_test_* to estimate PC scores from each method. Thus, even though the PeDecURe objective function involves different residualized versions of *X*, the features estimated using PC directions of variation from PeDecURe are based on the original *X*. This means that although PeDecURe is technically a supervised method (requiring that we have observed *Y* to compute residuals in the training sample), the directions of variation extracted from PeDecURe may generalizable to a testing sample where *Y* is unknown.

R code for implementing PeDecURe and for replicating the simulations below is available to reviewers upon request and will be made publicly available on GitHub upon acceptance for publication.

## 3 Parametric Simulation Studies

### 3.1 Simulation Setting

We first conduct a parametric simulation study to compare PeDecURe to PCA, Lin et al. (2016)’s AC-PCA, and PLS. Using the framework introduced above, our goal is to study associations between measured features (*X*) and a binary outcome (*Y*), while adjusting for a matrix of nuisance variables, *A*.

In each of 1000 simulations, we consider a sample size of *n* = 200 study participants and *r* = 300 observed features for each individual, which make up the rows of the *n* × *r* matrix *X*. We define *X* as a linear combination of *W* (an *n* × *p* matrix of the underlying patterns we are interested in, with *p* = 3) and *A* (an *n* × *q* matrix of nuisance variables, with *q* = 2). Our goal is to identify and estimate primary directions of variation, *v_j_* (*j* = 1, …, *p*) that correspond to the portion of *X* that is determined by *W*, but not by *A*.

In each simulation, for the *i*th subject and *j*th feature of interest,

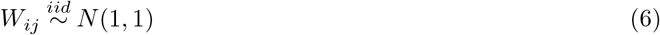

is the *i, j*th row and column entry of *W*, where *i* = 1, …, *n* and *j* = 1, …, *p* (with *n* = 200 and *p* = 3). The *i*th row of *W* will be denoted by *W_i_* below.

Next, we set *q* = 2 and randomly draw realizations of the subject-level nuisance variables from the following multivariate normal distribution:

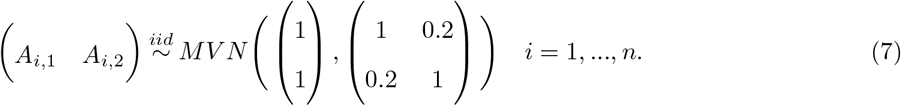

where 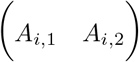 forms the *i*th row of the matrix *A*. Below, we will use *A*_1_ and *A*_2_ to denote the *n* × 1 vectors that make up the columns of *A*.

Next, the observed subject-level vector *X_i_* of *r* features, forming the *i*th row of *X*, is simulated as follows:

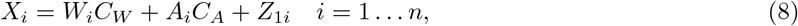

where *C_W_* and *C_A_* are matrices of constants specified below (with dimension *p* × *r* and *q* × *r*, respectively) and 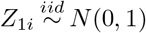. Following the notation described in Section 2.1, after combining the *X_i_* (*i* = 1, …, *n*) to form the rows of the population feature matrix, we center each column of *X* by its mean. (Note that, for consistency of notation, we will continue to use *X* to refer to the centered version of the simulated feature matrix.)

*C_W_* is defined such that row *k* (*k* = 1, …, *p*) has value *k* in positions (*k* – 1) × (*r/p*) + 1 through *k* × *r/p* and 0 elsewhere. For our case of *p* = 3 and *r* = 300, row *k* =1 has value 1 in positions 1 through 100; row *k* = 2 has value 2 in positions 101 through 200, and row *k* = 3 has value 3 in positions 201 through 300, while the remainder of the matrix is 0’s:

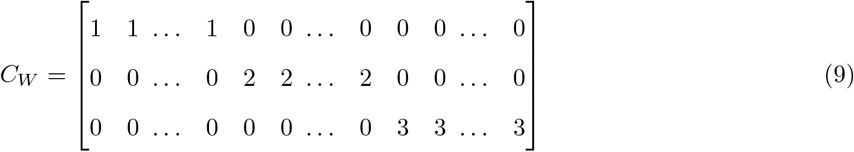

Similarly, *C_A_* is defined such that row *l* (*l* =1, …, *q*) has value *l* in positions (*l* – 1) × (*r/q*) + 1 through (*l* × *r/q*). For our case of *q* = 2 and *r* = 300, row *l* = 1 has value 1 in positions 1 through 150 and row *l* = 2 has value 2 in positions 151 through 300, and 0 elsewhere:

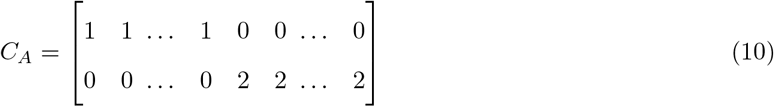

We then simulate the binary outcome variable *Y* as a function of the variables above. First, we compute subject-level probabilities (Pr(*Y_i_* = 1) = *P_Y_i__*) as follows (again in the case of *p* = 3 and *q* = 2):

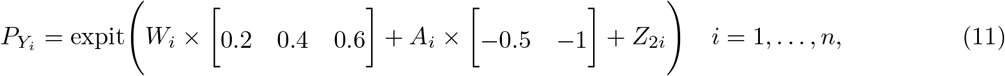

where expit() denotes the inverse logit function. Thus, each *P_Y_i__* is a linear combination of *W_i_* and **A*_i_* (each weighted by *p*- and *q*-dimensional vectors of constants, respectively), plus random noise 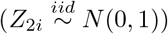. Finally, realizations of the binary outcome variable *Y_i_* are obtained through random draws from a Bernoulli distribution using the calculated probabilities:

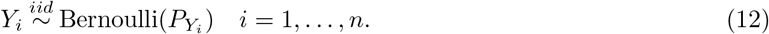

We apply PCA, AC-PCA, PLS, and PeDecURe to 70% of each simulated sample (*n_train_* = 140), estimating the top three primary directions of variation *v_j_*(*j* = 1, …, 3) from each method in this subset and reserving the remaining 30% of each simulated sample for testing (*n_test_* = 60). The tuning parameter λ for AC-PCA and PeDecURe is re-optimized in each training sample.

Partial Pearson correlation coefficients are then estimated between each PC score and *Y* (conditional on *A*_1_ and *A*_2_), *A*_1_ (conditional on *Y* and *A*_2_), and *A*_2_ (conditional on *Y* and *A*_1_). In evaluating the performance of each method in this parametric simulation setting, we will consider a method to perform well if the distribution of the absolute value of partial correlation coefficients between the PC1 score (in both training and testing samples) is higher than the distributions of these associations between all PC scores with *A*_1_ and *A*_2_.

### 3.2 Simulation Study Results

Overall, our parametric simulation results support the favorability of PeDecURe for estimating features that both preserve relatively strong partial correlations with a variable of interest while diluting those with nuisance variables.

Figure 2 shows that the distributions of partial correlations between the PC1 scores from PCA and PLS with *Y* (conditional on *A*_1_ and *A*_2_) appear slightly higher in 1000 simulated training (green outline, white boxes) and testing (green outline, grey boxes) samples compared with PeDecURe. However, the distributions of partial correlation coefficients between all three PC scores—especially the PC1 score—with *A*_2_ (orange outline) are high for both PCA and PLS, suggesting that the nuisance variables contribute to variation in the features estimated using these methods. PCA’s third PC score is also highly associated with *A*_1_ (purple outline), with partial correlations in both the training and testing sets exceeding 0.70. PC1 scores estimated using PeDecURe, on the other hand, have consistent and relatively higher distributions of partial correlations with *Y* (green outline) in both the training (white boxes) and testing (grey boxes) sample distributions. PC scores estimated by PeDecURe also have consistently lower distributions of partial correalations with each nuisance variable. Indeed, these distributions have higher variability in test samples, but they are nevertheless consistently lower relative to the distribution of partial correlations between the PC1 score from PeDecURe with *Y*.

**Figure 2:**
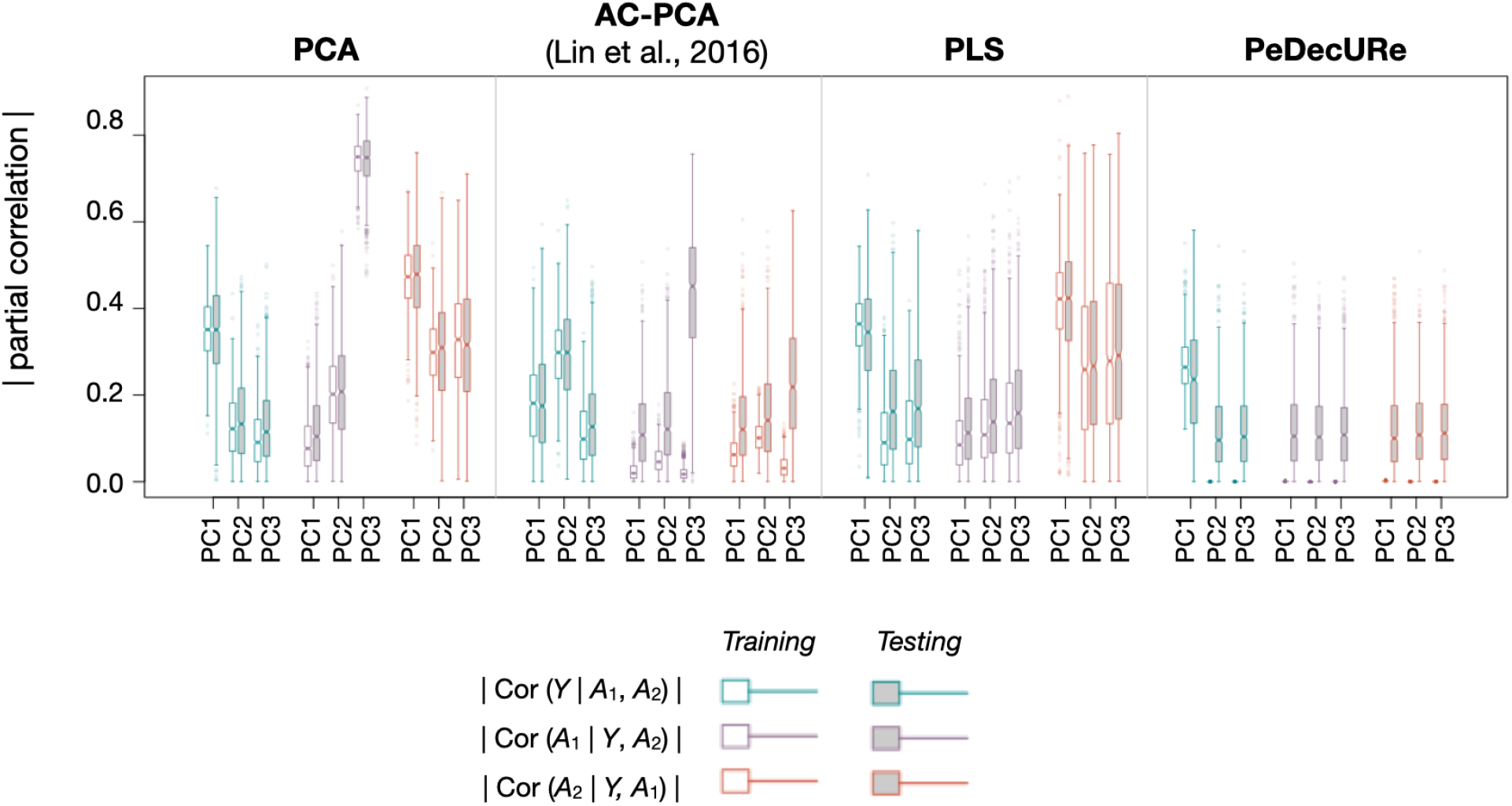
Simulation results in 1000 random training (white boxes) and testing (grey boxes) samples. Box border color indicates partial correlation with either the outcome variable (*Y*), or one of the two confounding variables (*A*_1_, *A*_2_), conditional on the other variables. Of the four methods considered, PeDecURe is the only method that produces a relatively higher distribution of absolute values of partial correlation coefficients between the first primary component (PC) score and *Y* (conditional on *A*_1_ and *A*_2_) and lower distributions of partial correlations between all three PCs and each nuisance variable.

Compared with Lin et al. (2016)’s AC-PCA, PeDecURe’s first PC score has slightly higher distributions of partial correlation coefficients with *Y* in training (white boxes) and testing (grey boxes) sets, and in the training set, PeDecURe’s distribution of partial correlations with *Y* is less variable. Although AC-PCA’s first two PC scores have relatively low distributions of partial correlation coefficients with both *A*_1_ (purple outline) and *A*_2_ (orange outline) in both training (white boxes) and testing (grey boxes) samples, the distributions of partial correlation coefficients between the third PC score and both nuisance variables in the test sample from AC-PCA is remarkably high.

## 4 Application to structural MRI data from the Alzheimer’s Disease Neuroimaging Initiative

### 4.1 Experiments using the ADNI MRI Data

Data used in the preparation of this article were obtained from the Alzheimer’s Disease Neuroimaging Initiative (ADNI) database (adni.loni.usc.edu). The ADNI was launched in 2003 as a public-private partnership, led by Principal Investigator Michael W. Weiner, MD. The primary goal of ADNI has been to test whether serial magnetic resonance imaging (MRI), positron emission tomography (PET), other biological markers, and clinical and neuropsychological assessment can be combined to measure the progression of mild cognitive impairment (MCI) and early Alzheimer’s disease (AD).

In this study, we apply PeDecURe, PCA, AC-PCA, and PLS to brain volumes extracted from baseline structural MRI scans from ADNI, linked with demographic variables and clinical phenotypes (ADNI team 2021) to evaluate prediction of and associations with a variable of interest (AD diagnosis, *Y*) and nuisance variables (age, *A*_1_ and sex, *A*_2_) using primary directions of variation estimated using each method. In practice, our goal is to identify a method that will enable us to quantify the direct relationship between ROI volumes in the brain, *X*, and AD diagnosis, *Y*. The relationship between cortical atrophy (as measured by ROI volumes) and AD has been examined over many years of AD research (Singh et al. 2006; Eskildsen et al. 2013). However, researchers have also documented significant differences in cortical atrophy rates and patterns across age and sex groups, including in numerous studies involving ADNI data (Double et al. 1996; Hua et al. 2010; Jiang et al. 2014). In this sense, both age and sex confound the relationship between brain structure and AD diagnosis, and controlling these sources of variation is a necessary intermediate step in any study seeking to directly relate AD diagnosis with patterns in the brain.

We pre-processed baseline structural MRI data from ADNI using Doshi et al. (2016)’s multi-atlas segmentation pipeline, estimating volumes for 137 regions of interest (ROIs) in *n* = 840 ADNI participants. Standardized baseline cognitive assessments were used to stratify participants into three diagnosis categories: At baseline, 230 participants were considered “cognitively normal” (CN), 200 met criteria for AD, and 410 were identified as having “mild cognitive impairment” (MCI). As an additional preprocessing step, we applied ComBat (Johnson et al. 2007; Fortin et al. 2017) to correct for batch effects associated with different scanner manufacturer (adjusting for diagnosis, sex, and age). Of the 840 participants in our subset, 447 were scanned in MRIs manufactured by GE, 58 Philips, 319 Siemens, and 16 were excluded because we were not able to identify their scanner manufacturers based on available data (Figure 3(a)). Finally, we restricted our sample to *n* = 422 participants who met criteria for CN and AD (*n_CN_* = 225 and *n_AD_* = 197 (Figure 3(b), after excluding those with MCI and those whose scanner manufacturer information was not available)).

**Figure 3:**
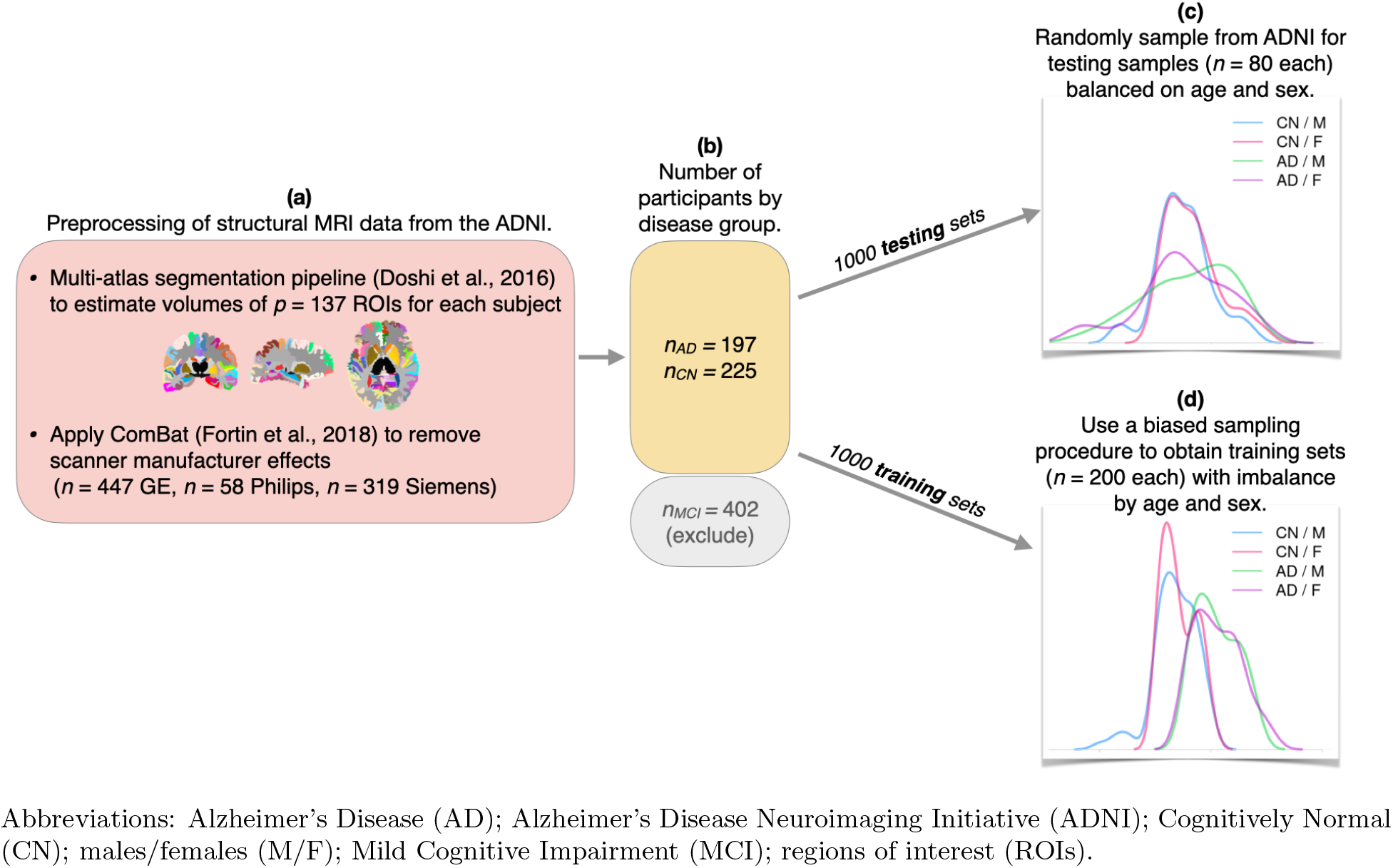
Preprocessing of ADNI data and sampling of training and testing samples for plasmode simulation study. Note: each of 1000 testing groups is sampled before the training sample, as it preserves the distribution of the original ADNI study. Each training sample is selected through a biased sampling procedure, which ensures that the age distribution for individuals with AD is higher than for those in the CN group, and females are more likely to have AD than males (see Section 4.1).

### 4.2 Plasmode Simulation Setting

Since models that ignore or inadequately control for nuisance sources of variation pose threats to generalizability, our simulation analysis of the ADNI data uses a flexible procedure to sample training and testing samples, such that the distribution of age and sex differs between these samples. Thus, our assessment of method performance in the testing sample should reflect each method’s generalizability in settings with shifted distributions of one or more nuisance variables.

We apply the following sampling procedure 1000 times to obtain different training and testing subsamples meeting certain criteria. We first set aside each testing sample, including *n* = 80 participants randomly subsetted from the 422 ADNI participants who met the inclusion criteria stated above in Section 4.1. Thus the typical testing sample preserves the same age and sex distributions across disease group that is present in the original ADNI data (Figure 3(c)). We then apply a biased sampling procedure to the remaining 342 participants (422 – 80), where we sample *n* = 200 individuals for each training sample using a biased sampling procedure to induce imbalance in both age and sex in each testing sample. Since older age is associated with both increased prevalence of AD and cortical atrophy (including in typical human aging) (Fjell et al. 2009), and since AD affects females at a higher rate than males (Laws et al. 2018), we use this sampling procedure so that the training samples would reflect an imbalanced distribution of age and sex not present in the original ADNI sample, but that would certainly be present (although to a lesser degree) in a random sample of adults. As illustrated in Figure 3(d), we restrict the age of AD participants to be 75 years or older, CN participants to be 80 years or younger, and set the probability of including a female with AD in a training sample about three times more likely than the probability of including a male with AD in the training sample.

We apply PCA, Lin et al. (2016)’s AC-PCA, PLS, and PeDecURe to each of the training samples to estimate the top three PC loadings from each method. We then evaluate each method’s performance in each training and testing sample after computing the top three PC scores in both samples. Similar to our assessment of our parametric simulations, we consider distributions of the absolute values of partial correlation coefficients between each PC score and *Y* conditional on *A*_1_ and *A*_2_, *A*_1_ conditional on *Y* and *A*_2_, and *A*_2_ conditional on *Y* and *A*_1_.

Next, we consider predictive performance using features estimated in the test samples using each method. Since by definition the first direction of variation estimated by each decomposition captures the largest percentage of variation (of some covariance matrix, which differs across methods), we assess prediction in the test sample of disease group, age, and sex using the PC1 score from each method alone. In practice, we will designate a method as having favorable performance if its PC1 yields high predictive accuracy for AD status (high area under the curve (AUC) for predicting *Y*) and poor prediction of both age (high root mean squared error (RMSE) for predicting *A*_1_, a continuous nuisance variable) and sex (low AUC or high 1–AUC for predicting *A*_2_, a binary nuisance variable). For each method, we fit simple logistic (for *Y* and *A*_2_) or linear (for *A*_1_) regression models with the training sample PC1 score as the sole independent variable. Estimated coefficients are then applied to the PC1 score from the test sample and AUCs (for *Y* and *A*_2_) or RMSEs (for *A*_1_) are calculated to assess out-of-sample prediction. We also compare distributions of these prediction metrics to those obtained from cross-validated models trained using the *n*=422 ADNI participants meeting inclusion criteria described above. Unlike the samples used to train the models used for prediction in our plasmode simulation setting, the sample of 422 ADNI participants has a balanced distribution of age and sex across disease groups.

Finally, since the PC loadings (e.g., *v*_1_, *v*_2_, *v*_3_) are each vectors with dimension *p* = 137, the number of ROIs, it is possible to interpret these loadings anatomically in the brain. For example, the estimated value in the *j*th position of *v*_1_ may be interpreted as a weight indicating the importance of that ROI (relative to the other ROIs) in explaining variation in the data. To identify the most important ROIs according to each decomposition method, we conduct an exploratory analysis where we identify the ROIs with estimated weights exceeding the 95th percentile of estimated weights (all the values in *v*_1_ from a given training sample and method). After identifying the top 5% of ROIs for each simulation, we identify which ROIs met these top 5% criteria for at least 50% of all simulations. Finally, we visualize and compare the top ROIs from the first direction of variation across methods. To facilitate interpretation in this exploratory analysis, we also identify the top 5% of ROIs in PC1 estimated by each method in the ADNI sample of *n*=422. Since the ADNI sample of *n*=422 participants has a balanced distribution of age and sex across disease groups, the top 5% of ROIs estimated in this sample may reflect what ROIs would be most important for explaining variation in the data in a setting that is, in theory, free from the nuisance variation that we induced into each training sample in our plasmode simulation setting.

### 4.3 Results from ADNI Plasmode Simulations

Distributions of the absolute values of partial correlation coefficients are plotted in Figure 4(a) and suggest that in both training and testing samples, PeDecURe performs as we intended it to. Specifically, associations between the PC1 score and diagnosis (*Y*, green outline) are highest for PeDecURe compared to PCA, AC-PCA, and PLS in both the training (white boxes) and testing (grey boxes) samples (although, as expected, there is higher variability in the partial correlation distributions in the test samples). Furthermore, PeDecURe’s scores on all of the first three PCs have low distributions of absolute partial correlation coefficients with both nuisance variables (age (*A*_1_, boxes with purple outline) and sex (*A*_2_, orange)), while PCA, AC-PCA, and PLS have noticeably higher distributions for at least one PC.

**Figure 4:**
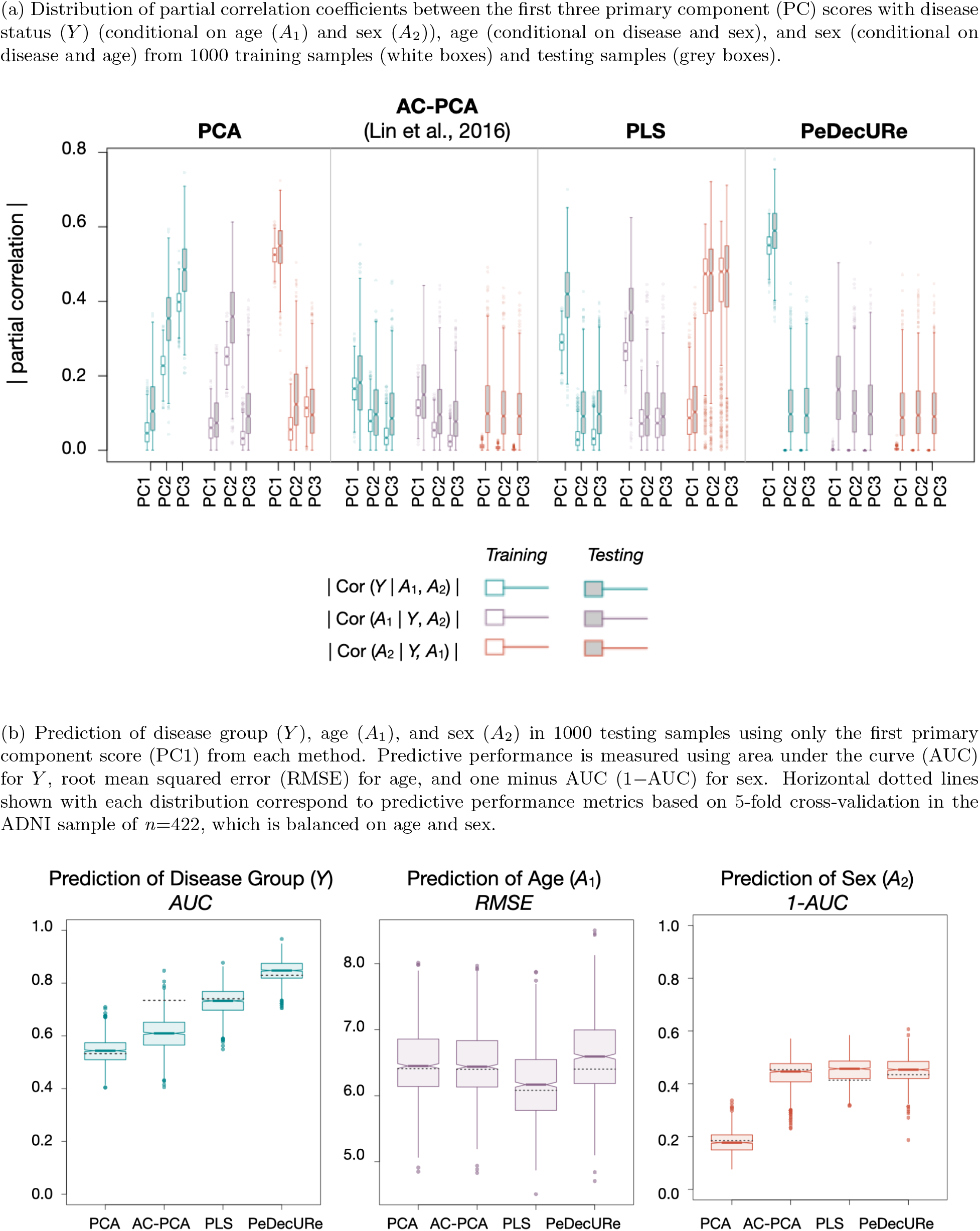
Plasmode simulation results using data from the Alzheimer’s Disease Neuroimaging Initiative.

As shown in Figure 4(b), PeDecURe outperforms the three other methods for disease group prediction. Not surprisingly, as PLS is the only other method that prioritizes maximizing covariance with the outcome (recall Figure 1), its PC1 score is also more highly predictive of diagnosis compared with PCA and AC-PCA. It is conceivable that the higher AUC distribution we obtain from a model involving PeDecURe’s PC1 score compared with a model involving PLS’s PC1 score may be due to the lack of nuisance variable adjustment in estimating PLS’s directions of variation.

It is still worth noting that PeDecURe has a similar RMSE distribution for age prediction as PCA and AC-PCA, and PeDecURe also has a similar 1–AUC distribution for sex prediction as AC-PCA and PLS. Therefore, in this plasmode simulation setting, PeDecURe’s favorability over the other methods is apparent in that its prediction of *Y* exceeds that of other methods and its prediction of confounding variables is no worse than other methods.

In Figure 4(b), horizontal dotted lines indicate the five-fold cross-validated prediction metrics according to an analysis of the full sub-sample of *n*=422 from ADNI included in this study, which has a balanced distribution of age and sex, so in theory we would not need to adjust for nuisance variables. Interestingly, the cross-validated AUC for disease prediction using AC-PCA’s PC1 is much higher than the distribution of AUCs from AC-PCA in the test sample. Although AC-PCA performs similarly to PeDecURe in its prediction of age and sex, this finding may suggest poor generalizability of AC-PCA.

### 4.4 Exploratory Multivariate Pattern Interpretation

Finally, we consider patterns extracted from each method by comparing the top ROIs from the 1000 plasmode simulation training samples to the top ROIs identified when each method was applied to the *n*=422 ADNI participant sub-sample whose age and sex distributions were balanced across disease groups. The top 5% of ROIs according to simulations in 1000 imbalanced training samples and those from the balanced testing sample are visualized in the left and right panels of Figure 5, respectively.

**Figure 5:**
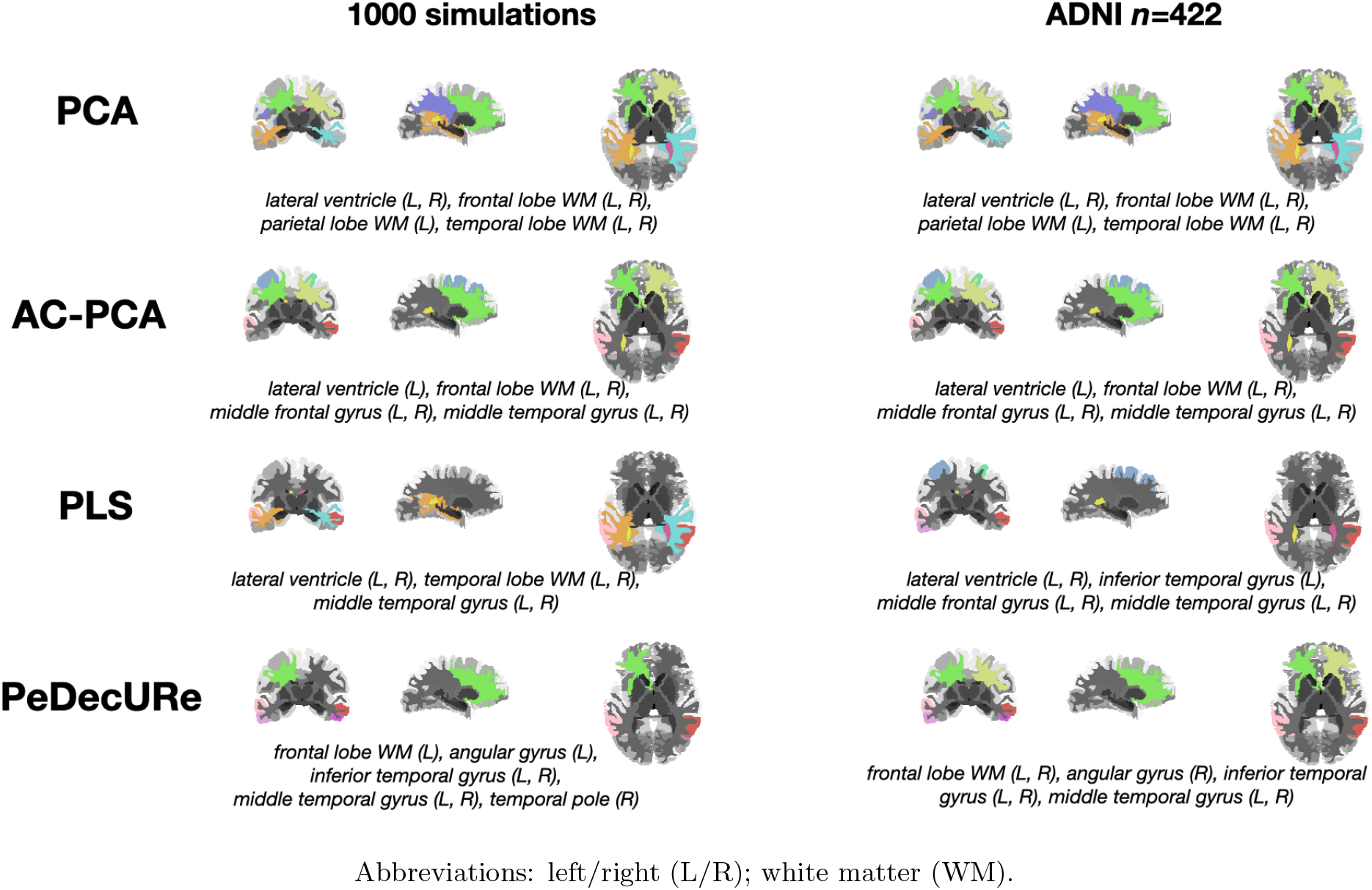
Top ROIs according to PC1 loadings from each method. ROIs above 95th percentile for at least 50% of simulations are shown on the left. On the right, we show the top 5% of ROIs after applying each method to the ADNI sample of *n*=422 (resulting group after applying exclusion criteria outlined in Figure 3).

It is interesting to note that the top 5% of ROIs identified by both methods that involve maximizing the covariance of *X* (PCA and AC-PCA) are the same in the imbalanced simulation and balanced analysis settings. Specifically, the same set of top ROIs we identify using PCA in the plasmode simulation setting and in the balanced ADNI sample of *n*=422 (Figure 5, left and right panels of first row are the same). The same is true for AC-PCA (Figure 5, left and right panels of second row are the same). We also note that the same is not true for PLS and PeDecURe (i.e., left and right panels differ in the last two rows of Figure 5), suggesting that feature extraction methods which prioritize shared variation in *Y* may be more sensitive to changes in the sampling distribution. This will be interesting to explore in greater detail in future work.

Since PeDecURe outperforms all other methods in disease prediction and having overall the lowest partial correlation distributions with nuisance variables (Figure 4), it seems plausible that PeDecURe may be best to identify the specific combination of ROIs that does best at balancing reduction in nuisance variation and preservation of disease associations. Nevertheless, across all methods, the top ROIs identified in both the plasmode simulations and analysis of the full ADNI sub-sample of *n*=422 are consistent with those implicated in previous structural neuroimaging-based studies of AD (Magnin et al. 2009; Habeck et al. 2008; Westman et al. 2011; Jiang et al. 2014). In future work, it will be interesting to develop non-parametric methods to quantify uncertainty in our interpretation of these ROIs.

## 5 Discussion

In this paper, we proposed Penalized Decomposition Using Residuals (PeDecURe), a method for estimating a low-dimensional set of features that have low correlations with nuisance variables but preserve strong associations with and are highly predictive of a variable of interest. PeDecURe combines strengths of PLS and Lin et al. (2016)’s confounder-adjusted principal component analysis (AC-PCA). Unlike these earlier methods, our method ensures that we prioritize only the portions of data that we care about (*X*\* residuals) and remove only the portions of the data that we do not want to preserve (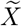 residuals).

While PeDecURe is motivated by the challenge of nuisance variable adjustment in MVPA of neuroimaging data, it adds to a growing body of methodological work (Linn et al. 2016; Rao et al. 2017; Adeli et al. 2019; Zhao et al. 2020; Aliverti et al. 2021). In the current paper, we focused on Lin et al. (2016)’s AC-PCA as a comparison to PeDecURe. AC-PCA has some important strengths, including that it is an unsupervised method (does not require that we observe *Y*) and can be used to correct for batch effects, with potential to outperform traditionally used batch effect correction methods, including ComBat and surrogate variable analysis (Johnson et al. 2007; Leek and Storey 2007). However, our parametric and plasmode simulation results suggested AC-PCA’s prioritization of nuisance variable removal differs from our goal, which is to reduce associations with confounders without also losing information about a variable of interest. In fact, we found that the incorporation of the outcome variable and residuals in PeDecURe also coincided with further reduction in nuisance variable associations compared with PeDecURe (Figures 2 and 4). Additionally, the comparatively poor prediction of *Y* and low partial correlations with *Y* using AC-PCA’s PC scores suggests features derived from this method may not be suitable for prediction. Our simulations also suggested that while PLS yields a PC1 score highly correlated with the outcome variable and AC-PCA produces PC scores with moderately lower partial correlations with confounding variables than PCA or PLS, neither PLS nor AC-PCA do well at both. In contrast, PeDecURe delivers on both, with consistently higher partial correlations between the PC1 score and outcome of interest, in both training and testing samples, and consistently low partial correlations between the first three PC scores and nuisance variables.

Like PLS and unlike AC-PCA and PCA, PeDecURe is indeed limited to training samples where the outcome (*Y*) is observed and, in this sense, would be classified as a supervised method. However, like PLS the directions of variation that we estimate have the flexibility of being applied in testing samples where *Y* is unobserved, since there is no need to residualize the test sample data, as we calculate PC scores using the original *X*, not residuals.

In this paper, we introduced a framework for estimating features which may reliably have reduced associations with nuisance variables and preserve associations with variables of interest, and we look forward to future extensions of this work. In particular, we hope to develop a collection of tools to facilitate interpretability of PeDecURe-derived features through nonparametric statistical tests, incorporate more flexible (e.g., nonlinear) models for residual estimation, test the performance of classical machine learning models trained using PeDecURe-derived features, extend PeDecURe to multivariate outcome variables, and extend PeDecURe to multimodal neuroimaging data.

## Acknowledgements

This work is supported by the following grants: R01MH112847 (RTS), R01MH123550 (RTS), R01NS112274 (RTS), R01NS060910 (RTS), U01AG068057 (CD), R01MH112070 (CD), and RF1AG054409 (CD). This work is also supported by the National Science Foundation Graduate Research Fellowship Program (SMW).

Data collection and sharing for this project was funded by the Alzheimer’s Disease Neuroimaging Initiative (ADNI) (National Institutes of Health Grant U01 AG024904) and DOD ADNI (Department of Defense award number W81XWH-12-2-0012). ADNI is funded by the National Institute on Aging, the National Institute of Biomedical Imaging and Bioengineering, and through generous contributions from the following: AbbVie, Alzheimer’s Association; Alzheimer’s Drug Discovery Foundation; Araclon Biotech; BioClinica, Inc.; Biogen; Bristol-Myers Squibb Company; CereSpir, Inc.; Cogstate; Eisai Inc.; Elan Pharmaceuticals, Inc.; Eli Lilly and Company; EuroImmun; F. Hoffmann-La Roche Ltd and its affiliated company Genentech, Inc.; Fujirebio; GE Healthcare; IXICO Ltd.; Janssen Alzheimer Immunotherapy Research & Development, LLC.; Johnson & Johnson Pharmaceutical Research & Development LLC.; Lumosity; Lundbeck; Merck & Co., Inc.; Meso Scale Diagnostics, LLC.; NeuroRx Research; Neurotrack Technologies; Novartis Pharmaceuticals Corporation; Pfizer Inc.; Piramal Imaging; Servier; Takeda Pharmaceutical Company; and Transition Therapeutics. The Canadian Institutes of Health Research is providing funds to support ADNI clinical sites in Canada. Private sector contributions are facilitated by the Foundation for the National Institutes of Health (www.fnih.org). The grantee organization is the Northern California Institute for Research and Education, and the study is coordinated by the Alzheimer’s Therapeutic Research Institute at the University of Southern California. ADNI data are disseminated by the Laboratory for Neuro Imaging at the University of Southern California.

